# CRISPRi screens reveal genes modulating yeast growth in lignocellulose hydrolysate

**DOI:** 10.1101/2020.09.17.301416

**Authors:** Friederike Gutmann, Cosimo Jann, Filipa Pereira, Andreas Johansson, Lars M. Steinmetz, Kiran R. Patil

## Abstract

**Background:** Baker’s yeast is a widely used eukaryotic cell factory, producing a diverse range of compounds including biofuels and fine chemicals. The use of lignocellulose as feedstock offers the opportunity to run these processes in an environmentally sustainable way. However, the required hydrolysis pretreatment of lignocellulosic material releases toxic compounds that hamper yeast growth and consequently productivity.

**Results:** Here, we employ CRISPR interference in *S. cerevisiae* to identify genes modulating fermentative growth in plant hydrolysate and in presence of lignocellulosic toxins. We find that at least one third of hydrolysate-associated gene functions are explained by effects of known toxic compounds, such as the decreased growth of *YAP1* or *HAA1*, or increased growth of *DOT6* knock-down strains in hydrolysate.

**Conclusion:** Our study confirms previously known genetic elements and uncovers new targets towards designing more robust yeast strains for the utilization of lignocellulose hydrolysate as sustainable feedstock, and, more broadly, paves the way for applying CRISPRi screens to improve industrial fermentation processes.

## Background

The baker’s yeast *S. cerevisiae* is the most frequently used eukaryotic cell factory (Berry et al., 2012; Ostergaard et al., 2000; Lopes et al., 2016 Jansen et al., 2017). The competitive production of biofuels and many other value-added compounds with yeast requires the use of cheap substrates that do not compete with food, feed and arable land. Lignocellulosic materials represent an economic and environmentally sustainable alternative feedstock. Spruce softwood is a promising lignocellulose source in the northern hemisphere (Willför & Holmbom, 2004; Zhu et al., 2009) and an abundant side product of the lumber, pulp and paper industry (Ragauskas et al., 2006). Lignocellulose has a complex structure, largely consisting of cellulose, hemicellulose and lignin. The extraction of fermentable mono-saccharide sugars (glucose) and hemicellulose polymers (pentose and hexose sugars) from cellulose biomass requires hydrolysis pretreatment. During this processing step, toxic compounds are released to the soluble hydrolysate which represent a major challenge in using this feedstock in an industrial setting (Pérez et al., 2002; Sun & Cheng, 2002; Almeida et al., 2007; Parawira and Tekere, 2011; Azhar et al., 2017). These compounds can be classified in three main groups: aliphatic acids, furan aldehydes and phenolic/aromatic derivatives (Jönsson et al., 2013). Aliphatic acids lower the intracellular pH and interfere with nutrient uptake (Pampulha and Loureiro-Dias, 1989). Furans inhibit dehydrogenases and raise levels of reactive oxygen species (Banerjee et al. 1981; Allen et al., 2010). Phenolics perturb plasma membrane composition and potential, resulting in a disruption of cell signalling and sorting processes (Keweloh et al., 1990). Understanding the impact of these substances on yeast growth and the cellular mechanisms of tolerance will enhance the use of lignocellulose-based biotechnological processes.

Several efforts have been made to characterize transcriptome changes (Sardi et al., 2016) and to improve substrate utilization and tolerance to lignocellulosic inhibitors in industrial yeast strains (Romaní et al., 2015, Cunha et al., 2018; Wu et al., 2017), including deletion collection screens for synthetic and straw lignocellulosic hydrolysates which linked tolerance mechanisms to ATPase activity and pH, pentose phosphate metabolism, lipid metabolism and the biosynthesis of amino acids (Pereira et al., 2014, Skerker et al., 2013). The traditional generation of deletion collections for industrial strains is laborious, since it requires the change of genomic sequence in multiple alleles, and cannot assess the effects of transcript down-regulation. The emerging CRISPR-based knock-out, interference and activation systems thus offer genetic screens with broader phenotypic scope.

Here, we established CRISPR interference (CRISPRi) screens to identify genetic functions that tune yeast growth in spruce hydrolysate. CRISPRi is emerging as a powerful tool to study genotype-phenotype relations via precise transcriptional repression (Gilbert et al., 2013 & 2014; Smith et al., 2016; McGlincy et al., 2020; Momen-Roknabadi et al., 2020). We employed an in-house developed single plasmid CRISPRi system (Smith et al., 2016) which is inducible by anhydrotetracyclin (ATc) (Farzadfard et al., 2013) to repress *S. cerevisiae* transcription factors (TFs, n=161) (De Boer and Hughes, 2012) and protein kinases (PKs, n=129) (Breitkreutz et al., 2010), key players in the regulation of cellular mechanisms and pathways. We detect genes capable of modulating yeast growth in hydrolysate, as well as in presence of toxic lignocellulose compounds to understand toxicity-dependent effects. We confirm previously known links to growth in hydrolysate and discover novel genetic associations that can be directly applied to advance sustainable bioprocesses (Fig. 1a).

**Figure 1.**
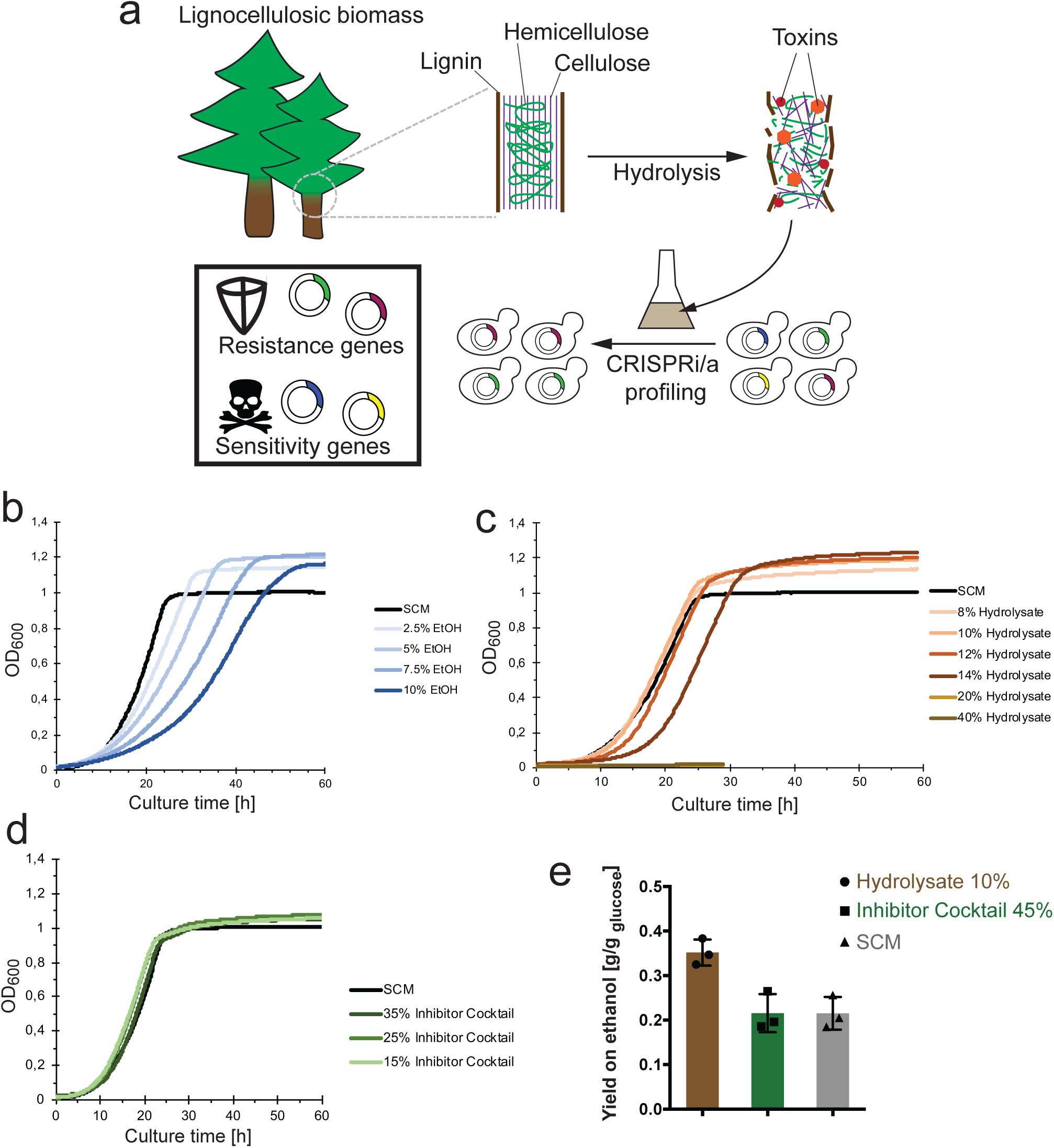
Study schematic and yeast tolerance to ethanol, hydrolysate and growth inhibitors. **(a)** Aim of the study. Schematic showing the hydrolysis of lignocellulosic material to convert large polymeric carbohydrates to mono-, di- and oligosaccharides, at the same time releasing toxic compounds that repel yeast growth (growth inhibitors). CRISPR interference or activation screens in hydrolysate allow for the identification of gene functions that contribute to stress sensitivity and resistance to enable the generation of robust strains for biotechnology applications. Schematic is inspired by Pérez et al. (2002) and Patel et al. (2016). Yeast growth in synthetic complete media with 2% glucose (SCM), in the presence of **(b)** ethanol, **(c)** spruce hydrolysate or **(d)** inhibitor cocktail with concentrations denoted, respectively. The optical density at 600 nm (OD600) (y-axis) of *S. cerevisiae* BY4743 strain cultures was measured in 96-well format over time (x-axis). Curves denote the average of n=4 wells, normalized by subtraction of media signal. **(e)** Ethanol yield obtained in different fermentation conditions is shown as g [EtOH produced] / g [glucose consumed], as calculated from HPLC measurements.

## Results

### Characterization of yeast fermentation in lignocellulosic hydrolysate

The diploid BY4743 yeast strain is able to grow in the presence of high amounts of ethanol (EtOH), with EtOH concentrations of 5% or 10% decreasing maximum growth rate by approximately 30% or 50%, respectively (Fig. 1b). We therefore decided to characterize hydrolysate tolerance in this strain which is genetically amenable and well characterized in contrast to process-specialised and frequently polyploid industrial strains.

We started by profiling growth and metabolite changes during fermentation in diverse conditions. Comparing growth in different dilutions of an industrially derived and widely used spruce hydrolysate (Demeke et al., 2014; Johansson et al., 2014; Koppram et al., 2012; Koppram et al., 2014; Soudham et al., 2014; Wu et al., 2017), we found that the BY4743 strain grew well in media supplemented with up to 14% hydrolysate without considerable changes in growth rate. However, in hydrolysate concentrations of 20% and more, no growth was observed (Fig. 1c). We further generated a mixture of eight selected growth inhibitors which are commonly found in lignocellulosic hydrolysate, based on previous studies (Tab. 1 and Methods). Growth profiles in presence of these compounds were similar to those in SCM, at least for the tested concentrations (Fig. 1d).

We then applied High Pressure Liquid Chromatography (HPLC) to quantify nutrient and metabolite changes during yeast cultivation in synthetic complete media (SCM), 10% hydrolysate and 45% inhibitor compound cocktail (IC). Remarkably, ethanol yields were increased by ∼65% in hydrolysate-supplemented medium compared to SCM and IC conditions (Fig. 1e). The increased ethanol production can partially be explained by the ∼25% higher initial glucose levels (Suppl. Tab. T1) and the break down of large sugar polymers during hydrolosis pretreatment that presumably increased the concentration of fermentable carbohydrates (Sardi et al., 2016), such as galactose and mannose (Francois et al., 2020; see Methods).

We further measured acetic acid in hydrolysate and IC conditions. The concentrations of this common growth inhibiting compound decreased by approximately 50% during fermentations (while glucose was fully consumed), suggesting that in these conditions yeast can metabolize acetic acid (Suppl. Fig. S1).

Taken together, we quantified dynamics in growth and metabolite abundances during fermentations in hydrolysate and in the presence of an lignocellulosic inhibitor mixture which showed that some growth-inhibitory substances can be metabolized and enabled us to derive optimal conditions for functional genomics assays.

### CRISPRi effects are reproducible and capture positive controls

To identify genes that modulate yeast growth in the presence of lignocellulose hydrolysate and inhibitor cocktail, we employed a plasmid-based CRISPRi system that is inducible with Anhydrotetracycline (ATc) to set up gene dosage screens. Each condition was assayed in triplicate in CRISPRi-inducing (+ATc) as well as in reference (-ATc) conditions (Fig. 2a). Multidimensional scaling (MDS) of read count samples indicate high similarity between replicates and allow to estimate sources of variability of CRISPRi effect size (Fig. 2b). Samples of the inhibitor cocktail were positioned closely to hydrolysate samples, suggesting higher similarity to this condition than compared with SCM (Fig. 2b). Selection was performed for ∼25 generations for all screens (Fig. 2a) to enable direct comparison of effects on doubling time. For a detailed comparison of fitness effects on multiple layers (Fig. 2a), we provide correlations of read counts (Suppl. Fig. S2), guide RNA (Suppl. Fig. S3) and gene log2 fold changes (log2FCs) across all screens (Suppl. Fig. S4).

**Figure 2.**
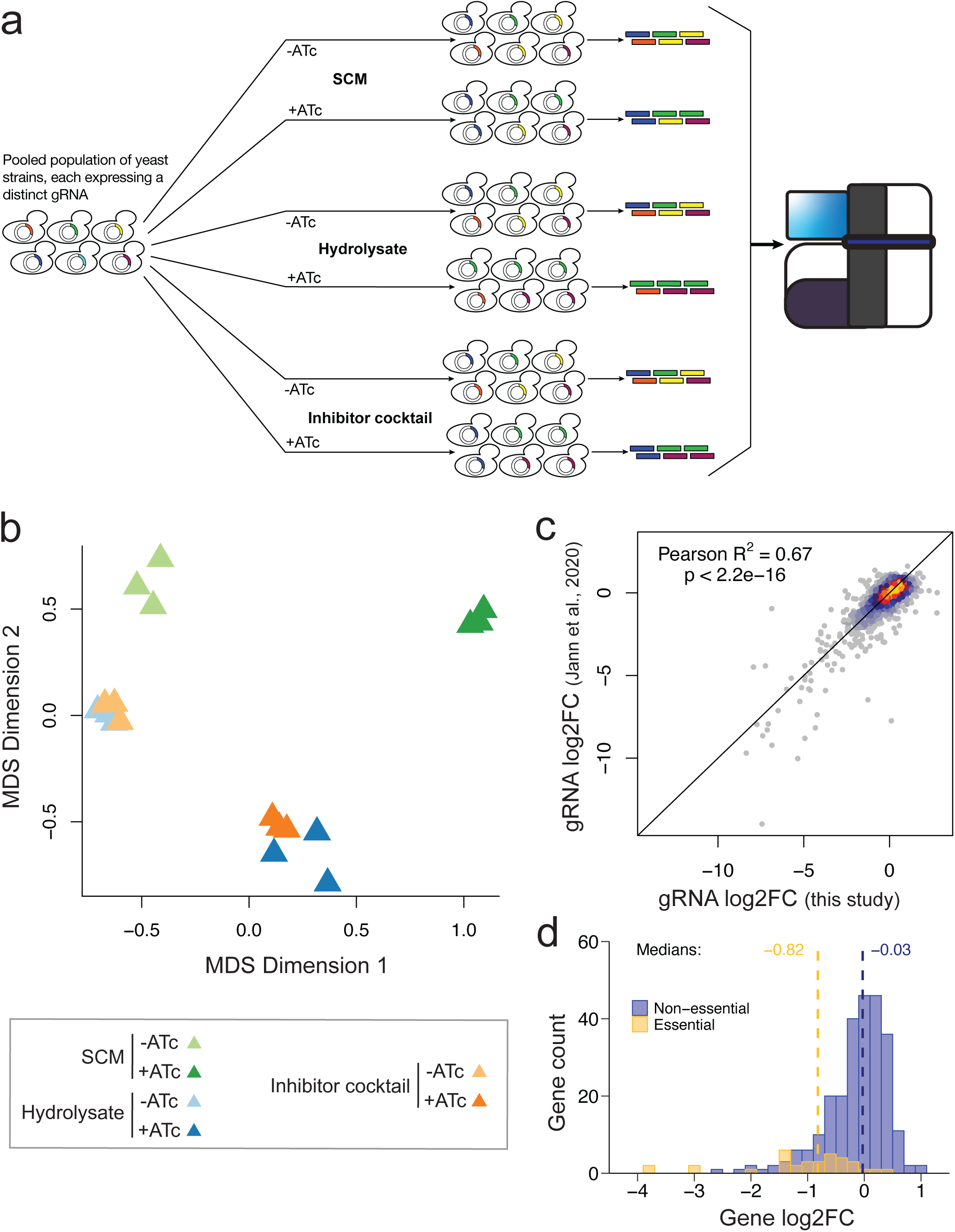
CRISPRi screens are reproducible and capture positive controls. **(a)** Schematic of screen procedure and selection conditions. All selections are performed with populations grown with ATc to induce CRISPRi and without ATc as reference, and in biological triplicate cultures. **(b)** MDS-plot of read count samples, depicting Euclidian distance variation in two dimensions (x- and y-axis) to estimate (dis)similarity of replicate samples and effect size between CRISPRi-induced (+ATc) and reference samples (-ATc) across all screen conditions. **(c)** Correlation of guide log2FCs from this study to a previous screen from Jann et al., (2020). CRISPRi effects of both screens report on fitness in SC medium. Jann et al. (2020) phenotyped the TF and PK libraries separately with two replicates while in the presented screens we phenotyped a single consisting of a combined TF and PK libraries and measured triplicates. **(d)** Distribution of log2 gene fold changes for non-essential (blue) and essential genes (gold).

Guide RNA log2FCs are highly correlated with those of a previous study where the TF and PK libraries have been profiled individually with different transformation batches and without oxygen limitation (Jann et al., 2020), indicating high reproducibility of CRISPRi effects (Pearson R^2^=0.67, p-value<2.2e-16, Fig. 2c). Accordingly, 67% of variation between CRISPRi effects observed in one of the screens were explained by the other. Interestingly, we measured increased fitness for the repression of three genes (*HAP1, RIM11* & *RME1*) that were not significant in the previous study (Jann et al., 2020), presumably due to the oxygen-limited conditions applied here to focus on fermentation. This is supported by the heme activated protein 1 (Hap1) TF which represses Rox1 in non-stress conditions (Zitomer and Lowry, 1992). In response to hypoxia, Hap1 is inhibited to de-repress Rox1 and induce hypoxic stress gene expression (Keng, 1992; Kwast et al., 1998; Zitomer and Lowry, 1992) which would be enhanced in CRISPRi strains. The Rme1 TF and the Rim11 PK (one of four glutathione synthetase kinase 3 homologs) regulate meiosis, and their knock-down effects may hint to growth-antagonizing roles in oxygen-limited environments. As anticipated from a competitive fitness assay, genes essential for viability (based on the *Saccharomyces* Genome Database SGD; Cherry et al., 1998), showed higher depletion compared to non-essential genes and represent positive controls that validate the experimental setup (Fig. 2d).

### Hydrolysate-specific gene functions and their applications in biotechnology

Having validated gene dosage effects in SCM and their reliability across studies, we screened CRISPRi populations in 10% spruce hydrolysate (Fig. 2a). This revealed fitness effects (gene fold change FDR<0.05 and at least two gRNAs with absolute log2FC>=1 and FDR<0.05) for the repression of 24 genes (Fig. 3a and Suppl. Tab. T2). Ten of these also showed growth effects in SC media (brown dots in Fig. 3a). Four more genes also caused general fitness defects although missing the strict significance requirements in the SCM screen. This left ten genes with hydrolysate-associated roles and minor or no impact on growth (labelled in Fig. 3a). Repression of seven genes decreased (*YAP1, HAA1, HOG1, PBS2, UGA3, CDC15* and *UME6)* and of three genes increased growth (*DOT6, SKO1* and *BUB1*) specifically in hydrolysate. Most of these have well-known roles in adaptation to diverse kinds of stress. Repression of *YAP1* results in decreased growth in SCM+10% hydrolysate medium (Fig. 3a). The Yap1 TF induces gene expression in response to oxidative stress (Kluge et al., 1997; Okazaki et al., 2007) and is known to be activated by toxic compounds including furans (Kim and Hahn, 2013) and phenolic molecules (Nguyen et al., 2014). In line with CRISPRi effects, the *yap1Δ* mutant has decreased fitness in hydrolysate (Skerker et al., 2013), and a recent study also demonstrated that *YAP1* overexpression increases growth in spruce hydrolysate (Wu et al., 2017), thus providing a direct validation of screen effects and illustrating how gene functions can be applied for strain design in biotechnology.

**Figure 3.**
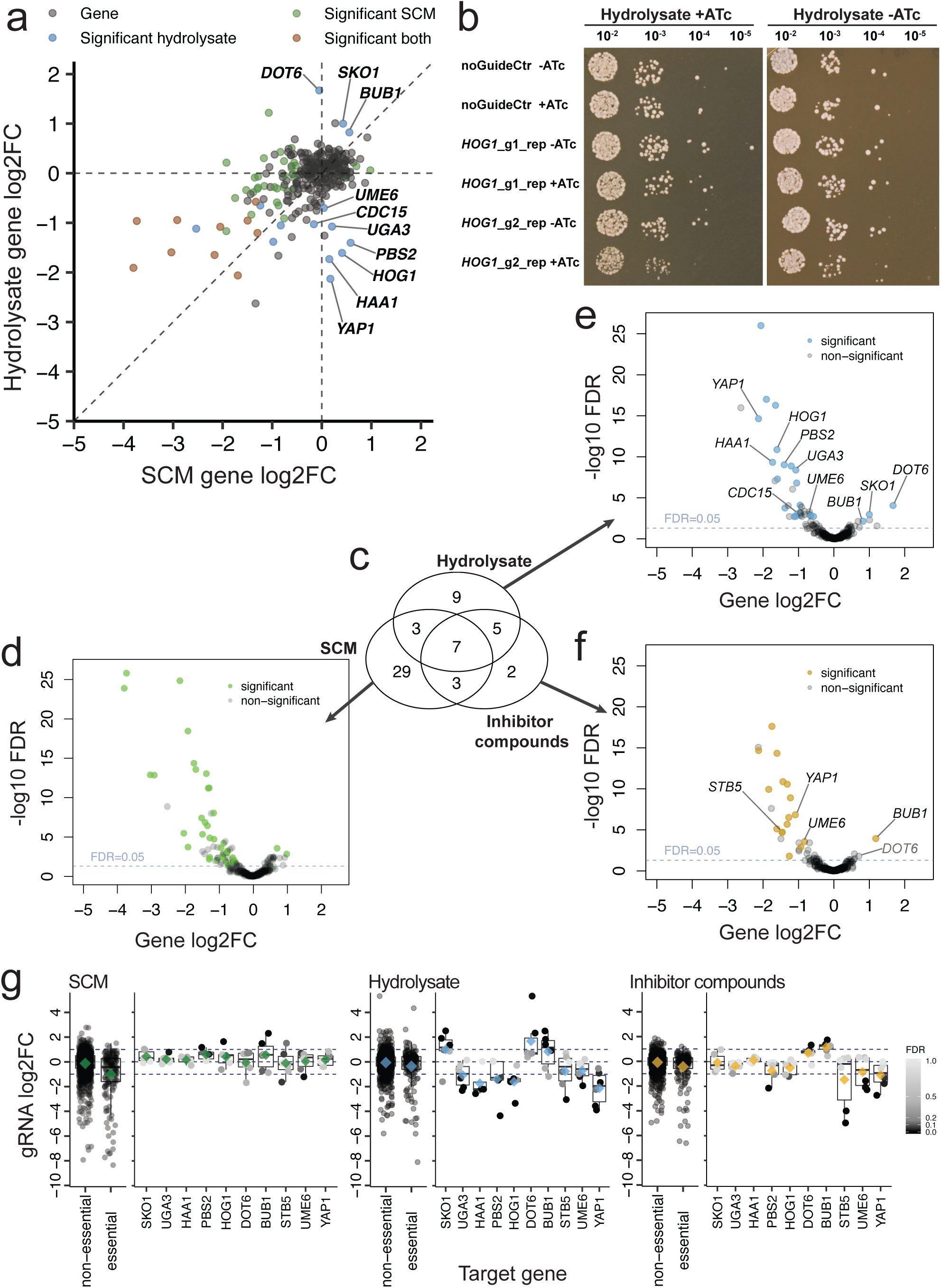
Genes with specific functions in hydrolysate fitness. **(a)** Scatter of gene log2FCs in SCM versus hydrolysate conditions. Dots denote essential (brown) and non-essential genes (green). Genes with strong hydrolysate-specific effect on fitness are labelled. **(b)** Dilution spot plating of Hog1 CRISPRi strains grown with or without 250ng/µl ATc for 24h and plated on SCM+10% hydrolysate agar plates with or without ATc. Cultures of Hog1 CRISPRi strains pre-grown in repressed (+ATc) and reference conditions (-ATc) are plated. Control strains harbor non-functional gRNAs. **(c)** Venn diagram of genes modulating fitness in SCM, hydrolysate and inhibitor cocktail conditions. The overlap of significant genes is shown together with volcano plots to illustrate perturbation strength and confidence of individual hits. Volcano plot of gene log2FCs and -log10 Benjamini Hochberg FDRs for CRISPRi effects in **(d)** SCM, **(e)** hydrolysate, and **(f)** in presence of inhibitor compounds. The dashed blue line marks an FDR of 0.05. Some but not all significant modulators are labelled for clarity. **(g)** Guide RNA log2FCs for selected genes across conditions. Dots denote gRNA log2FCs and are coloured by FDR for single genes (not for non-essential and essential gene panels). Diamonds denote the means and are coloured in green for SCM, blue for hydrolysate and yellow for inhibitor compound conditions akin to colours used in d-f. For genes, the mean gRNA log2FCs (diamonds) correspond to their gene log2FC.

Likewise, we measured decreased fitness in hydrolysate for *HAA1* in CRISPRi screening (Fig. 3a). The Haa1 TF is required for adaptation to mildly acidic environments (Fernandez et al., 2005), so that effects are most likely explained by the acidic pH of spruce hydrolysate (pH 4.5 of the 10% hydrolysate medium, compared to pH 5.5 of SCM). Akin to *YAP1*, the overexpression of *HAA1* in an industrial yeast strain increased growth rate in hydrolysate and additionally improved ethanol production (Cunha et al., 2018).

We further measured decreased fitness in hydrolysate in strains repressing *UME6* (Fig. 3a), which encodes a regulator of meiotic and translation-related genes (Lardenois et al., 2015). Accordingly, the *ume6Δ* mutant is sensitive to oxidative (Brown et al., 2006), hyperosmotic (Dudley et al., 2005) high temperature stress (Jann et al., 2020) and to more than 20 diverse chemicals some of which may have properties similar to lignocellulosic compounds (based on SGD; Cherry et al., 1998).

We further measured decreased growth in hydrolysate upon repression of *PBS2* and *HOG1* (Fig. 3a). The two mitogen-activated protein kinases (MAPKs) are central components of the high osmolarity glycerol (HOG) pathway which mediates adaptation to hyperosmotic environments. We confirmed effects on *HOG1* by dilution spot plating of individual CRISPRi strains expressing two gRNAs with moderate and strong effect size, respectively (Fig. 3b). In line with these results, the *hog1*Δ deletion mutant is also reported with decreased hydrolysate fitness (Wagner et al., 2019). HOG pathway effects could be due to the high osmolarity of spruce hydrolysate as a result of dissolved compatible solutes which are released from lignocellulosic material during hydrolysis, including salts and carbohydrates. In contrast, repression of the HOG-downstream TF *SKO1* increased cell growth in the presence of hydrolysate. Sko1 is constitutively nuclear and bound to promoters for repression in non-stress condition (Proft & Serrano, 1999; Pascual-Ahuir et al., 2001; Rep et al., 2001). Upon hyperosmotic stress which likely prevails in hydrolysate, Hog1 is activated and transitions into the nucleus where it phosphorylates Sko1 which is then exported to the cytosol, activating transcription via de-repression (Proft & Serrano, 1999; Pascual-Ahuir et al., 2001; Rep et al., 2001). Interestingly, genes induced in lignocellulose hydrolysate are enriched for Sko1 target genes (Sardi et al., 2016) which supports the relevance of Sko1-controlled transcripts in hydrolysate.

The mechanisms underlying further hydrolysate-specific effects of *BUB1, CDC15* and *UGA3* could not linked to their known functions and offer leads for future investigation (Fig. 3a). We notably observed increased growth in hydrolysate for repression of *DOT6* (Fig. 3a) which encodes a transcriptional repressor that responds to osmotic and oxidative stress (Lippman and Broach, 2009, Pascual-Ahuir et al., 2001). The stimulated fitness in hydrolysate upon *DOT6* repression can, akin to *SKO1*, be explained through de-repression. Proteins encoded by genes that affect growth in hydrolysate form physical interaction networks with bundled interactions at components of the osmotic (Hog1, Pbs2, Sko1) and oxidative stress response (Yap1, Dot6) and connect to growth regulators (Suppl. Fig. S5)

To better understand the cellular processes remodelled in hydrolysate, we determined the target genes of TFs that modulated fitness in hydrolysate using chromatin immunoprecipitation (ChIP) data (Sadowski et al., 2013) which were enriched for functions in translation, RNA binding, cyclic compound binding, the regulation of carbon metabolism (glycolysis and gluconeogenesis) and nuclear export of non-coding RNAs (Suppl. Fig. S6). We further searched for the phosphorylation targets of PKs modulating hydrolysate fitness, and found that these were enriched for functions in general kinase activity, Mitogen Activated Protein Kinase (MAPK) and target of rapamycin (TOR) signalling, mitophagy, stress granule components, chronical cell aging, as well as the cellular responses to osmotic stress, organic substances and acidic chemicals (Suppl. Fig. S7). Toxic chemicals, low pH and hyperosmotic conditions thus represent major stressors that yeast cells genetically brace against during growth in spruce hydrolysate.

### Contributions of lignocellulosic growth-inhibitors

We next screened for growth effects in the presence of lignocellulosic inhibitor compounds to determine genetic effects linked to their toxicity. After 25 generations selection in SCM+45% Inhibitor Cocktail, we identified seven genes with inhibitor-specific functions compared to SCM, five of which overlap with hydrolysate-specific effects (Fig. 3c, Suppl. Tab. T2). Fitness effects observed for repression of *HAA1, HOG1, PBS2, SKO1* and *UGA3* in hydrolysate were not measured in presence of inhibitor compounds and are therefore likely independent of the toxicity caused by the used substances and concentrations (Fig. 3g). In contrast, the strong hydrolysate-specific growth effects of *BUB1, DOT6, UME6* and *YAP1* CRISPRi strains were reproduced with the inhibitor cocktail and were thus caused by one or multiple compounds in the cocktail (Fig. 3g). In addition, we find that the repression of *STB5* reduced growth in inhibitor-supplemented and hydrolysate media although the gene barely missed the strict significance requirements in the hydrolysate screen (Fig. 3g). *STB5* encodes a zinc cluster activator of pleiotropic multidrug resistance genes (Akache et al., 2004). While *stb5Δ* strains are reported with decreased hydrolysate growth (Skerker et al., 2013), its overexpression was shown to increase growth in spruce hydrolysate (Wu et al., 2017). Three genes with inhibitor-specific effects are transcribed from bidirectional promoters which are hard to dissect by CRISPRi (*CKA2*|*SLD7, STE7*|*DHH1, FKH2*|*YNL067W-A*) since effects can be caused by the perturbation of either of the two genes, or their combined impact. Two of these loci (*CKA2*|*SLD7, STE7*|*DHH1*) were also significant in the hydrolysate screen. Clustering of gene fold changes across screens allows for the identification of specific and shared gene functions between conditions (Suppl. Fig. S8).

## Discussion

Here, we employed CRISPR interference screens to identify regulatory genes capable of adjusting the growth of baker’s yeast in spruce hydrolysate and in the presence of lignocellulosic toxins. This allowed us to explain contributions of toxicity in hydrolysate growth conditions, and how genetic screens can be utilized towards overcoming current challenges in hydrolysate fermentation.

CRISPRi perturbations are powerful to probe genotype-phenotype relations in high throughput, with low cost and labour, and with high reproducibility. Our single plasmid CRISPRi system has been deeply characterized (Smith et al., 2016 & 2017; Jaffe et al., 2019; Jann et al., 2020), is freely available on addgene (#73796) and supported by an in-house developed gRNA design platform (http://lp2.github.io/yeast-crispri/; Smith et al., 2016) and a customizable computational analysis pipeline (Methods). The plasmid system can be transformed into any strain background, including polyploid and industrial strains, to devise strategies for improving process performance.

The presented CRISPRi screens on yeast growth in hydrolysate complement genetic screens with deletion mutants (Pereira et al., 2014, Skerker et al., 2013). We found that hydrolysate-specific functions are frequently connected to stress adaptation in the responses to oxidative (Yap1, Stb5), osmotic (Hog1, Pbs2, Sko1), acidic (Haa1) and general stress (Dot6, Ume6). These mirror the environmental conditions prevailing in the hydrolysate which are presumably perceived as stressful by yeast cells. Overcoming them can partially be tackled by the pre-treatment of hydrolysate by, e.g. de-salting, pH adjustment or by addition of reducing agents to quench reactive oxygen species. Alternatively, or in addition, yeast strains can be genetically engineered to enhance growth, stress tolerance and productivity. Here we demonstrate that CRISPRi screens provide the opportunity to identify gene targets for strain optimization and, notably, multiple of our hits were already known to affect yeast growth in hydrolysate and have been successfully applied, including Yap1, Stb5 and Haa1. Overexpression of these three genes has been shown to increase tolerance to hydrolysate (Cunha et al., 2018; Wu et al., 2017) which not only validates their genetic functions but also demonstrates their potential to improve bioprocesses.

Our screens suggest novel genes, expression of which can be modified to optimize hydrolysate fermentation. These include decreasing expression levels of *BUB1, DOT6* and *SKO1*; and overexpressing *UGA3, UME6*, and the HOG pathway components *PBS2* and *HOG1*. While our aim was to characterize genetics underlying fermentative growth, employing the screen under other selection conditions, such as low pH (Koivistoinen et al., 2013), high ethanol (Fujita et al., 2006), and high temperatures (Banat et al., 1998; Edgardo et al., 2008; Hasunuma and Kondo, 2012; Jann et al., 2020; Van Vleet and Jeffries, 2009), could be used to further improve specific processes.

Since growth-inhibiting agents are described as major challenge for utilizing plant hydrolysates for fermentation, we screened for fitness effects in presence of eight such compounds covering aliphatic acids, furan aldehydes and phenolic/aromatic molecules (Jöonsson et al., 2013). While our cocktail was defined to largely cover these prevalent impacts, there are presumably additional lignocellulosic substances with yet other mechanisms of toxicity. Our finding that at least one third of hydrolysate-specific gene functions were explained by toxicity effects therefore represents a lower bound and likely an underestimation. Due to the high prevalence of toxicity effects, relieving them through genetic regulation is an attractive avenue to facilitate the fermentation of lignocellulosic material.

## Conclusion

Taken together, we show how CRISPRi screens can be used to identify genetic elements underlying complex environmental conditions encountered by cells in industrial bioprocesses. Our study pinpoints genetic functions that can be engineered to facilitate utilization of lignocellulose as feedstock for yeast fermentation, and thereby hopefully motivates and contributes to the establishment of environmentally sustainable procedures in biotechnology.

## Methods

### Yeast strains, bacterial strains and plasmids

Two gRNA libraries targeting sets of 129 protein kinases (Breitkreutz et al., 2010) with 668 gRNAs and 161 transcription factors (De Boer and Hughes, 2012) with 885 gRNAs were used as described in Jann et al. (2020). Libraries were integrated in the dCas9-Mxi1 plasmid system (Smith et al., 2016), transformed into the BY4743 strain background and pooled together. All chemical compounds, oligonucleotides, plasmids, as well as bacterial and yeast strains used in this study are listed in Additional File AF1.

### Growth media

Filter-sterilized synthetic complete uracil-dropout medium (SCM-Ura) containing 20 g/L glucose, 6.7 g/L yeast nitrogen base without amino acids and with ammonium sulfate and 2 g/L amino acids uracil-drop-out mix, pH 5.5, was used for fermentation. SCM-Ura was supplemented with hydrolysate or inhibitor cocktail mixture as indicated. Spruce hydrolysate was kindly provided by SEKAB (Örnsköldsvik, Sweden). The sawdust raw material contains 13 g/kg dry solid/dry weight (DS) arabinose, 20 g/kg DS galactose, 408 g/kg DS glucose, 50 g/kg DS xylose, 108 g/kg DS mannose, 0.26% ash (525°C), 2.3% cyclohexane/acetone soluble matter and 27 % DS lignin (Klason method) as reported by SEKAB. The sawdust with a dry matter content of 50% was pretreated with sulfur dioxide, enzymatically hydrolysed and filtered to remove solids. The resulting hydrolysate is of defined composition and has been chemically characterized with 83 g/l glucose and higher order carbohydrates (26 g/l mannose, 9 g/l xylose, and less than 4 g/l galactose and arabinose), as well as toxic compounds that include, among others, 4.7 g/l acetic acid, 3.4 g/l 5-hydroxymethyl-furfural, 1.2 g/l furfural and less than 1 g/l of phenolic derivatives, levulinic and formic acid (Alriksson et al., 2011). In CRISPRi screening, SCM media was supplemented with 10 % of described hydrolysate, pH 4.5.

We defined an inhibitor cocktail based on previous characterizations of spruce and other plant hydrolysates, focussing particularly on medium produced by SEKAB (Demeke et al., 2014; Johansson et al., 2014; Koppram et al., 2012; Koppram et al., 2014; Soudham et al.; 2014; Wu et al., 2017) or generated under similar technical and chemical conditions (Larsson et al., 1999; Martín et al., 2018; Petersson & Lidén, 2007). Inhibitor compounds were purchased from Sigma Aldrich, and mixed to a 5x concentrated cocktail before addition to SCM (Table 1). Coniferyl aldehyde, vanillin and p-Hydroxybenzaldehyde were diluted in 3 ml 0.1 M NaOH prior to addition. Inhibitor-supplemented SC-Ura was adjusted to pH 5 with 0.1 M NaOH.

**Table 1.**
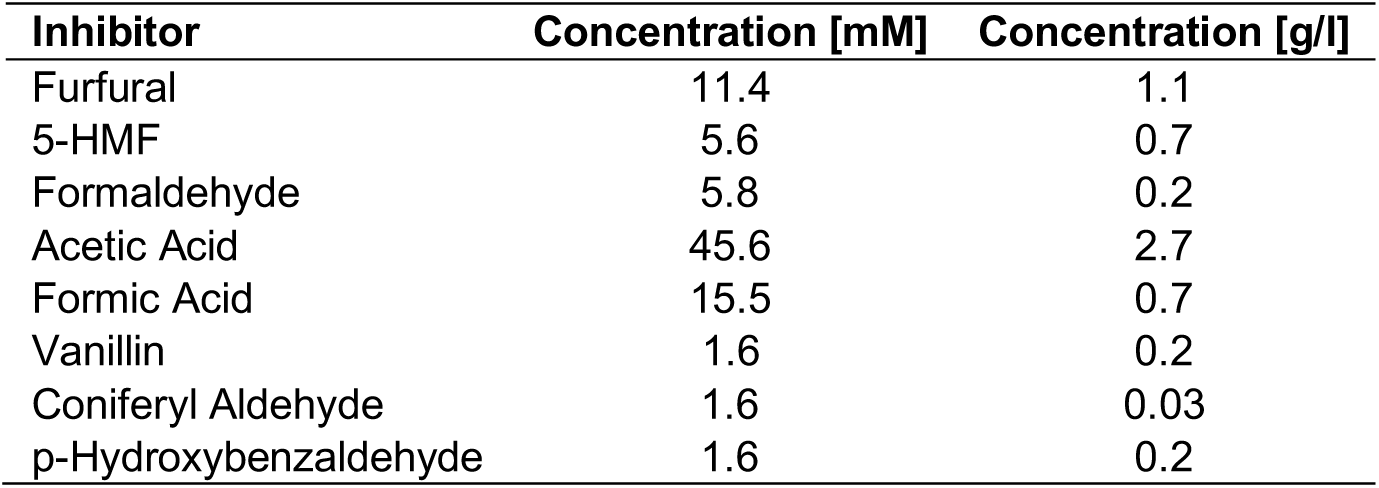
Composition of toxic compounds of the inhibitor cocktail. Compound concentrations are listed in mM and g/l resembling the final (1x) concentration in SCM.

### Analytical methods

Quantitative determination of acetate, ethanol and glucose was performed by High-performance liquid chromatography (HPLC). The HPLC system was equipped with a Refractive Index Detector (Alliance HPLC with 2414 RID, Waters, Eschborn, Germany) and a Rezex ROA-Organic Acid H+ (8%) column (Phenomenex, Aschaffenburg, Germany) which was held at 65 °C during all measurements. 0.5 mM Sulfuric acid was used as mobile phase carrying the samples at a flow rate of 0.5 ml/min. Samples were stored at 6 °C prior to each run which took 15 min. Hydrolysate containing samples were diluted four times. Compounds were quantified by comparing the metabolite peak in the sample with a mixture of standards with known concentrations of each metabolite.

### Dilution spot plating

CRISPRi cultures were grown in SCM + 10% hydrolysate either with or without 250 ng/ml ATc for 24h at 30°C. A dilution series was prepared in medium without ATc, and 10µl of the dilutions were plated on SCM + 10% hydrolysate agar plates without or with 250 ng/ml ATc (added to agar-containing medium after autoclaving). Photographs were taken after 2 days incubation at 30°C to report on colony size.

### Culture conditions

For HPLC experiments, yeast cultures were grown in 50ml in shake flasks at 30°C with 175 rpm. Flasks were sealed with rubber plugs to mimic oxygen-limiting conditions, stimulating fermentation rather than respiration. Plugs were pierced with sterile needles (27, BD Microlance, Becton Dickinson) and cotton plugs to enable CO2 exchange. Plate reader assays were performed under oxygen-limited conditions in 96-well plates (Nunclon Delta Surface, Thermo Scientifc) with Synergy HTX Multi-Detection Microplate Readers (BioTek) at 30°C. Plates were inoculated with an optical density at wavelength 600 nm (OD600) of 0.005. OD600 was measured in 15 min intervals and shaken at 800 rpm for 10 s before measurements. For screens, yeast was cultured in sealed falcon tubes as described below.

### Competitive growth assays

For screens, defrosted hydrolysate and inhibitor cocktail were diluted in SCM-Ura to 10% and 45%, respectively. Screens were performed in triplicate samples, each with addition of 250 ng/ml ATc to induce gRNA expression and without ATc as reference. Yeast cultures were profiled in 15 ml falcon tubes containing 11 ml medium that was inoculated with overnight cultures at OD 0.005. Falcon tubes were sealed to mimic oxygen-limited conditions, and a needle (20 G, BD Microlance) stuffed with cotton was pierced through the lid. Cells were then grown in a shaking incubator at 30 °C and 180 rpm. The tubes were opened only to assess growth stage. Cells were transferred to fresh medium in late mid-exponential phase (before reaching OD600=1). During the screen, two such transfers were performed to maintain exponential growth. Cell samples were centrifuged, and pellets were washed and used to extract plasmid DNA.

### Sequencing

Plasmid DNA was purified using the Miniprep kit (QIAprep Spin, Qiagen, Hilden, Germany) with modified protocol. Cell pellets were resuspended in P1 solution (accordingly to kit mannufacturer) and incubated with with 9U lyticase (Sigma-Aldrich) at 37 °C for 30 minutes, followed by harsh vortexing for 2 minutes. The remaining steps were performed following the provided kit protocol. Illumina sequencing adapters and inline barcodes were introduced to DNA barcodes by PCR, creating a specific double index of samples. All PCR products were analysized by gel electrophoresis and then purified (QIAquick, Quiagen). Samples were pooled to similar amounts. The resulting sequencing library was concentrated by performing another PCR purification, and a 1% agarose gel electrophoresis was performed for size-selection and purification. DNA quality was checked with a DNA-Bioanalyzer, and a diluted sample was sequenced on an Illumina Next-Seq500 machine in paired-end mode with 75 bases read length.

### Data processing and analysis

Raw Illumina sequencing reads were demultiplexed using Jemultiplexer. Trimmed reads were aligned to a gRNA sequence reference genome with Burrows-Wheeler aligner to compute read counts. Read counts were processed in R code with the edgeR package and our computational pipeline allows to perform this with alternative read count-based analysis package, such as limma or DESeq2. Significant genes were required to have a gene log2FC with FDR<0.05 and at least two supporting gRNAs with FDR<0.05 and absolute log2FC>=1.

### Enrichment analyses

Gene ontology-enrichment was performed using the gProfiler2 R package (Reimand et al., 2019). TF target genes were determined using Chromatin Immuno-Precipitation on chip (ChIP-chip) data from Gonçalves et al. (2017). Genes bound by two or more TFs from the set of significant modulators identified in CRISPRi screens were used for enrichment analysis with the *S. cerevisiae* genome as statistic background. Phosphorylation targets of protein kinases were determined with data from phosphogrid 2.0 (Sadowski et al., 2013). The PK targets of significant modulators of CRISPRi screens were used for enrichment analysis, using all protein kinase targets measured in the phosphoproteomics data set as statistic background.

### Data visualization

Plots were generated in R (V. 3.4.1)(R Core Team, 2018) with the ggplot2 (V. 3.1.0) (Ginestret, 2011), ggally (V. 1.3) (Schloerke et al., 2017) and pheatmap (V. 1.0.10) packages (Kolde, 2012). In boxplots, the middle line denotes the median, and lower and upper hinges denote the first and third quartiles, respectively. Figures were designed in Adobe Illustrator 2019.

## Supporting information

Reagents, strains and oligos

gRNA fold changes

Gene fold changes

## Abbreviations

ATc: Anhydrotetracycline
CRISPRi: Clustered Regularly Interspaced Short Palindromic Repeats interference
(d)Cas9: (dead) Clustered Regularly Interspaced Short Palindromic Repeats-associated protein 9
HPLC: High performance liquid chromatography
IC: Inhibitor compound cocktail
MDS: Multidimensional scaling
SCM: Synthetic complete medium

## Availability of data and materials

Demultiplexed sequencing data, read counts, gRNA fold changes and gene fold changes were deposited at Gene Expression Omnibus and are accessible under GSE155590 with the token *chkxawwunlgrjeb*. Computed gRNA barcode and gene level fold changes are provided in the supplementary as Additional File AF2 and AF3, respectively.

## Contributions

C.J., F.G., F.P. and K.R.P. conceived the study. C.J., F.G. and F.P. designed experiments. C.J. and F.G. performed experiments and data analysis. A.J. and L.M.S. provided essential insight and advice. C.J., F.P. and K.R.P. supervised the study. C.J. wrote the manuscript with help from F.G., F.P. and K.R.P. All authors reviewed the manuscript and approved the final version.

## Acknowledgements

The authors wish to thank Antonius J.A. van Maris (Royal Institute of Technology, Stockholm, Sweden) for his supervisory support of the Master’s Thesis of F.G. Adnan Cavka from SEKAB (Örnsköldsvik, Sweden) is gratefully acknowledged for the kind provision of pretreated lignocellulose hydrolysate and technical advice. The authors very much appreciated the useful discussions with Maurizio Bettiga (Chalmers University of Technology, Gothenburg, Sweden) regarding the experimental setup and Justin Smith (Stanford University) for CRISPRi construct setup.

## Supplementary Information

## Supplementary Figures

**Supplementary Figure S1.**
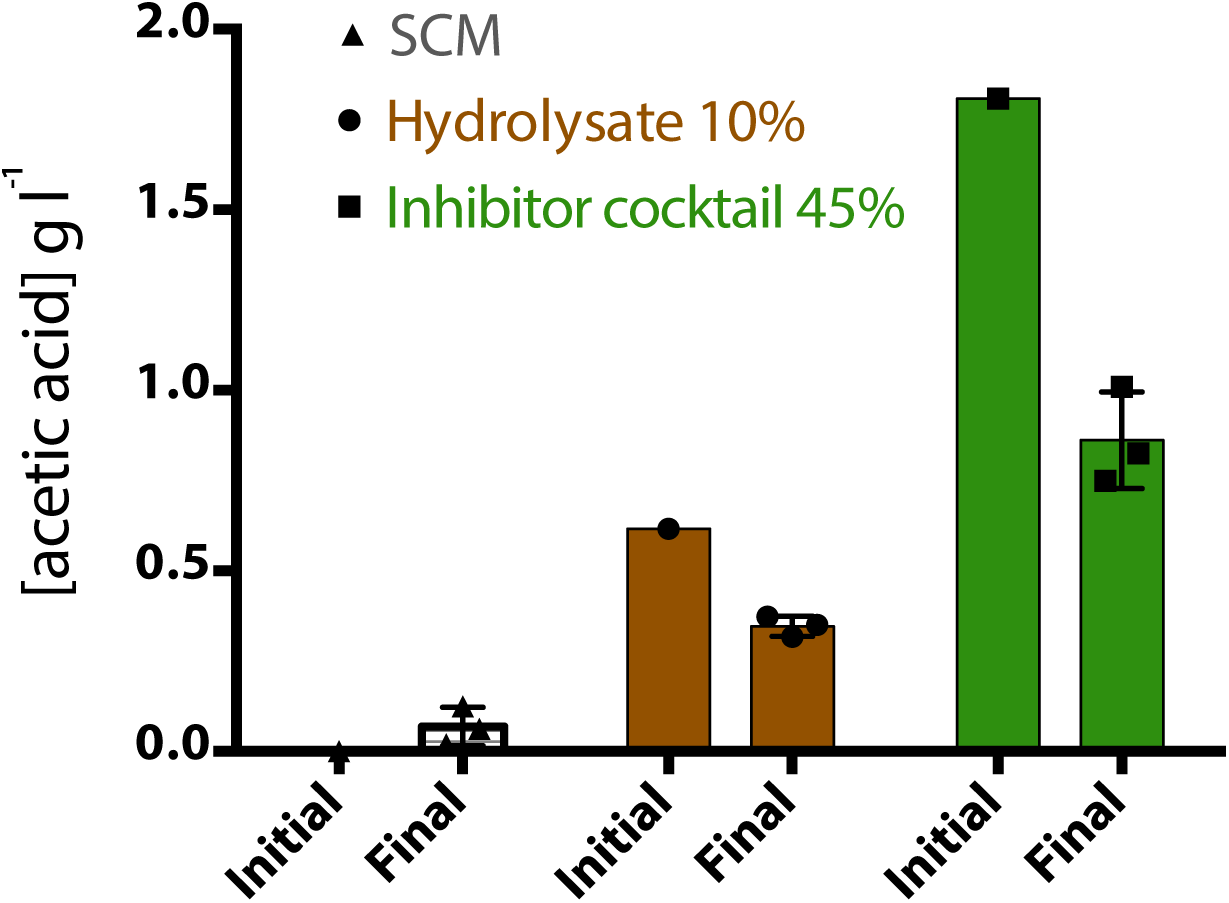
Acetic acid metabolization. Changes of acetic acid (in g/l) during fermentation in different growth conditions (indicated in figure legend) at cultivation start and end points, measured by HPLC.

**Supplementary Figure S2.**
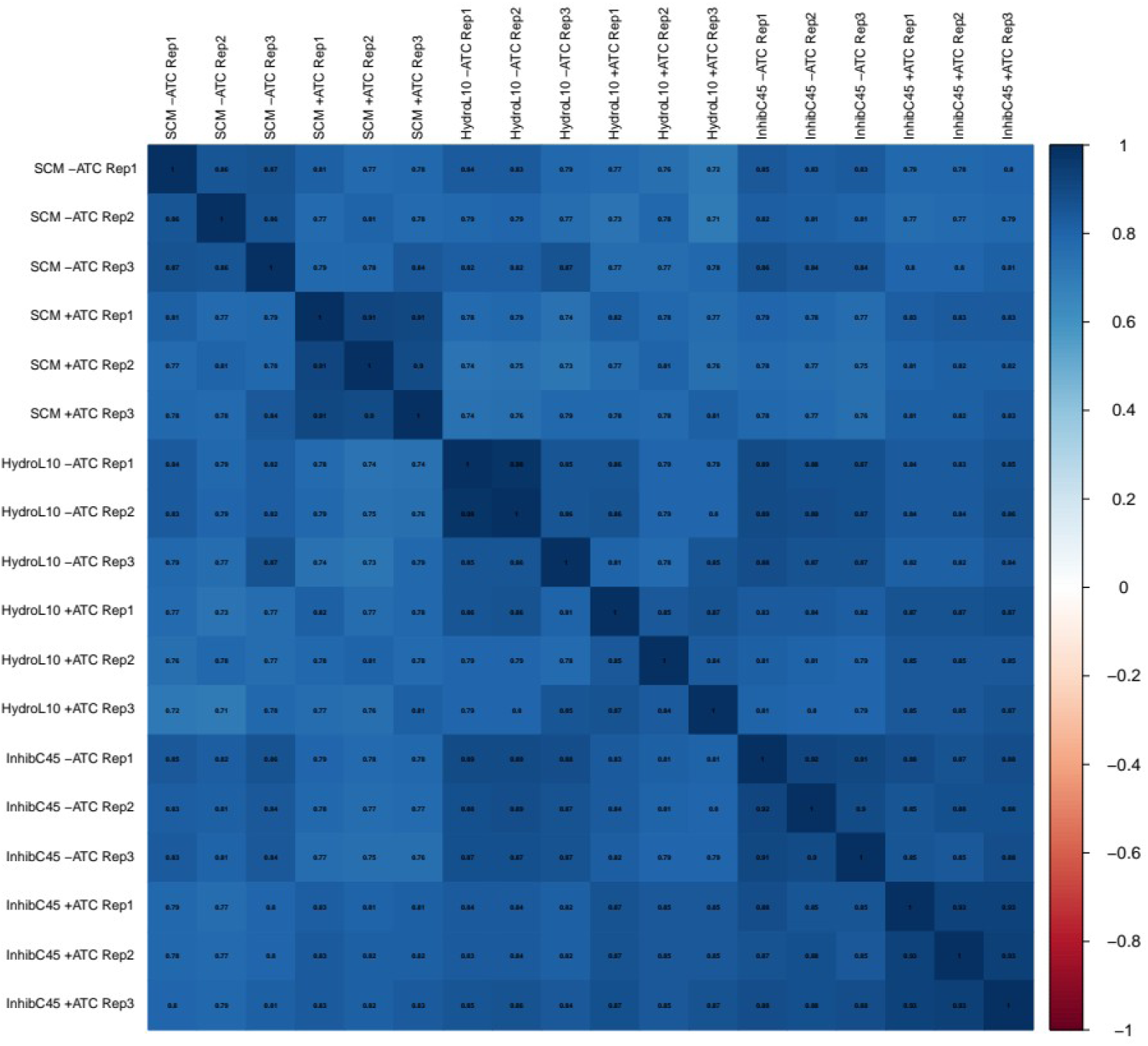
Read count correlation. Spearman correlations of read count samples across screens.

**Supplementary Figure S3.**
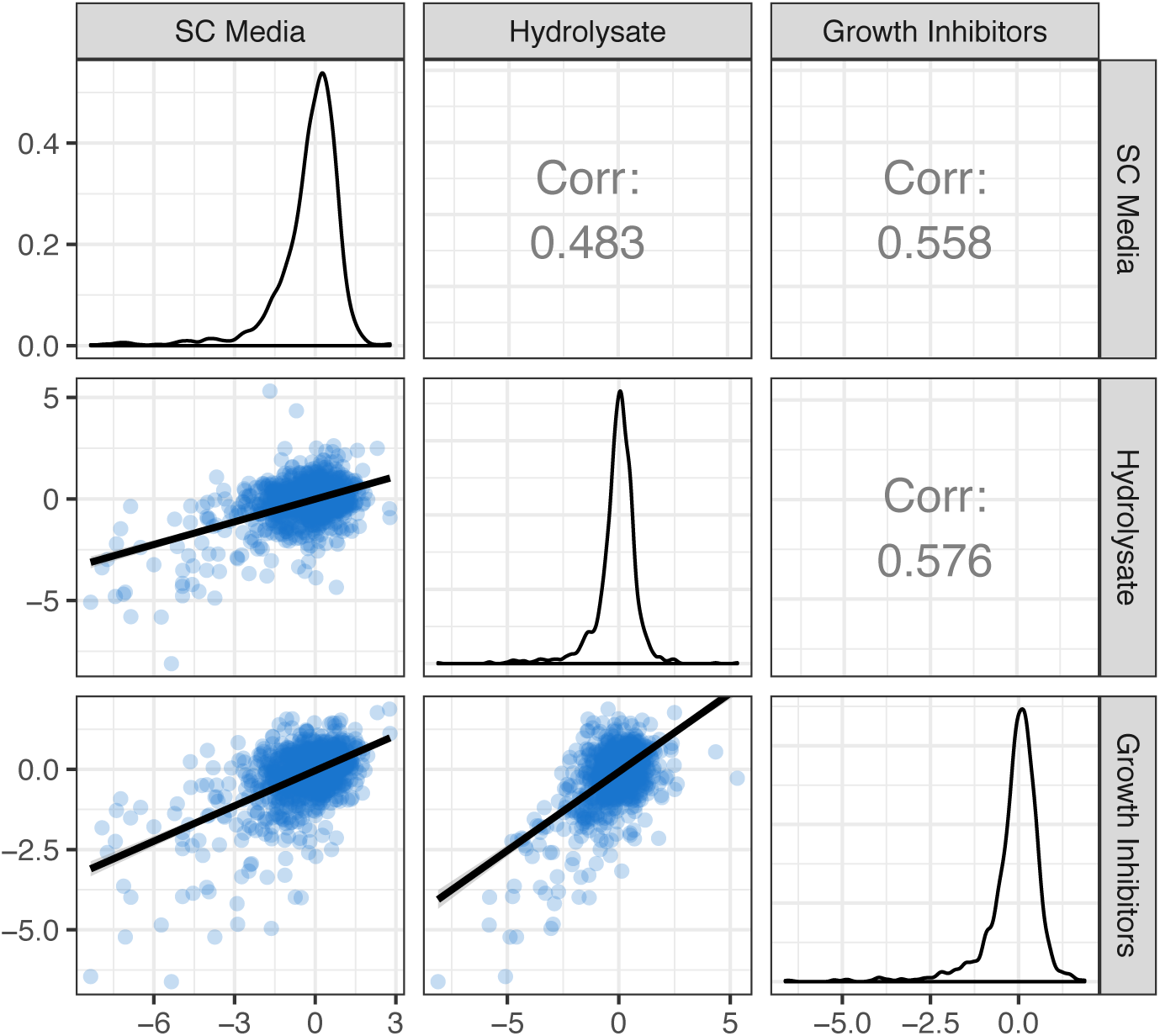
Guide RNA fold changes across conditions. Scatter plots with dots denoting gRNAs, density distributions and Pearson correlations of gRNA log2 fold changes across screen conditions.

**Supplementary Figure S4.**
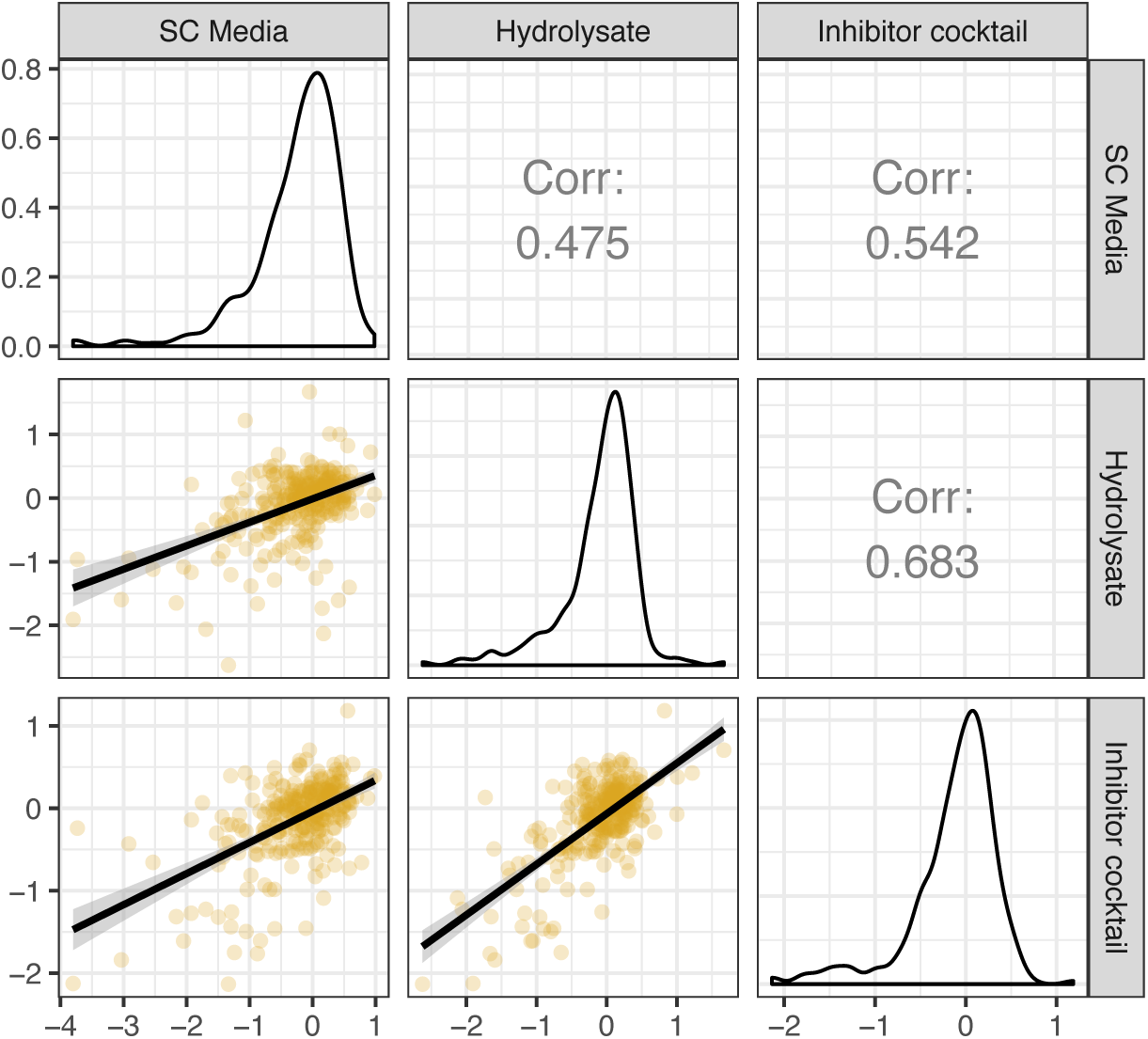
Gene fold changes across conditions. Scatter plots with dots denoting genes, density distributions and Pearson correlations of gene log2 fold changes across screen conditions. Line denotes smoothed linear fits.

**Supplementary Figure S5.**
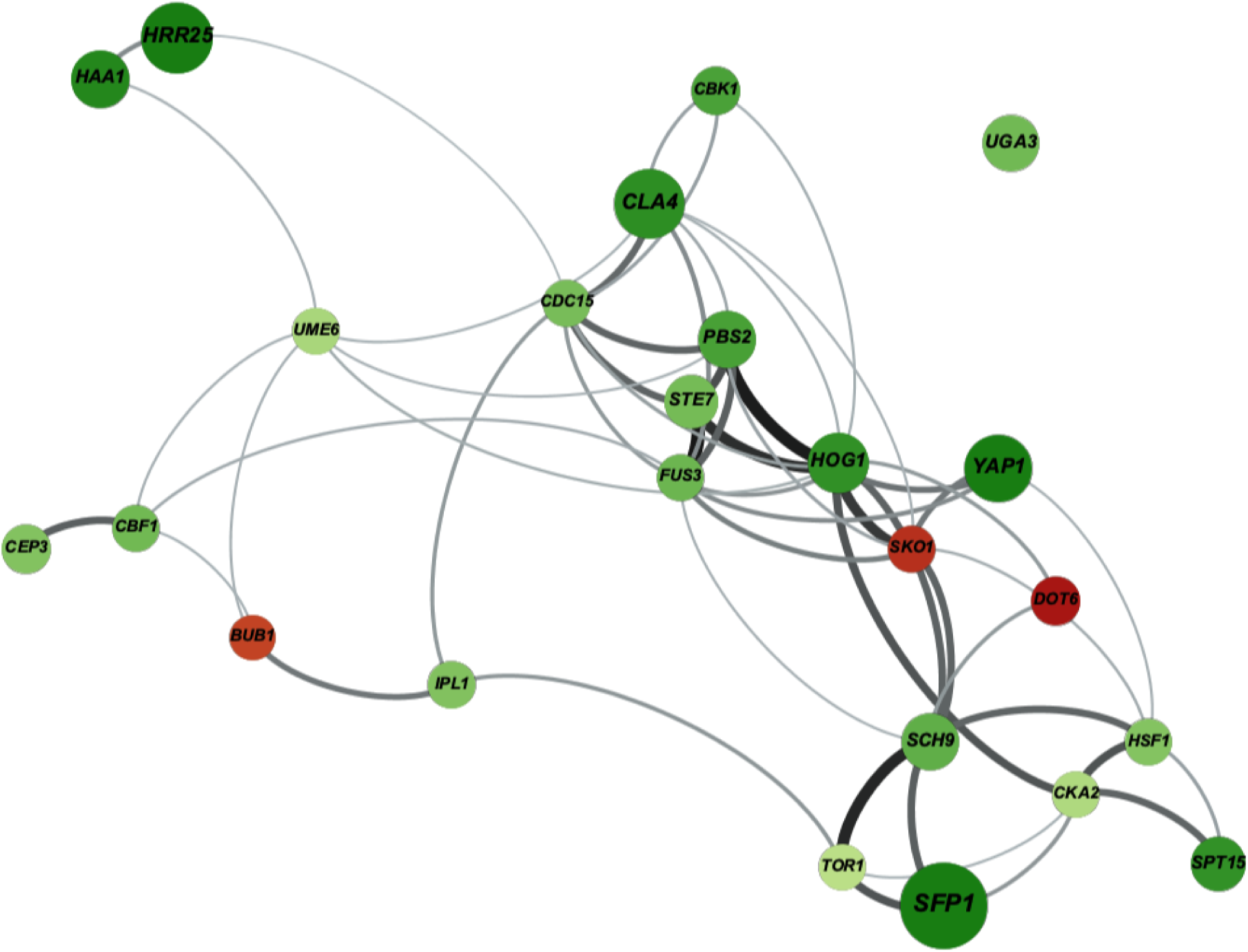
Protein-protein interaction network between modulators of hydrolysate growth. Experimental protein-protein interactions of genes modulating cellular fitness in hydrolysate, obtained from STRING (Snel et al., 2000). Dots denote genes, coloured by gradients from light to dark by increased strength in either positively (green) or negatively (red) modulating hydrolysate fitness, obtained from screen log2 fold changes. Dot and gene label size denote the multiple-testing adjusted FDRs obtained in the screen. Line thickness indicates confidence of the physical interaction obtained from the STRING database. Network visualization was performed with Gephi (Bastian et al., 2009), using the Force Atlas 2 algorithm for clustering with standard parameters (Jacomy et al., 2012).

**Supplementary Figure S6.**
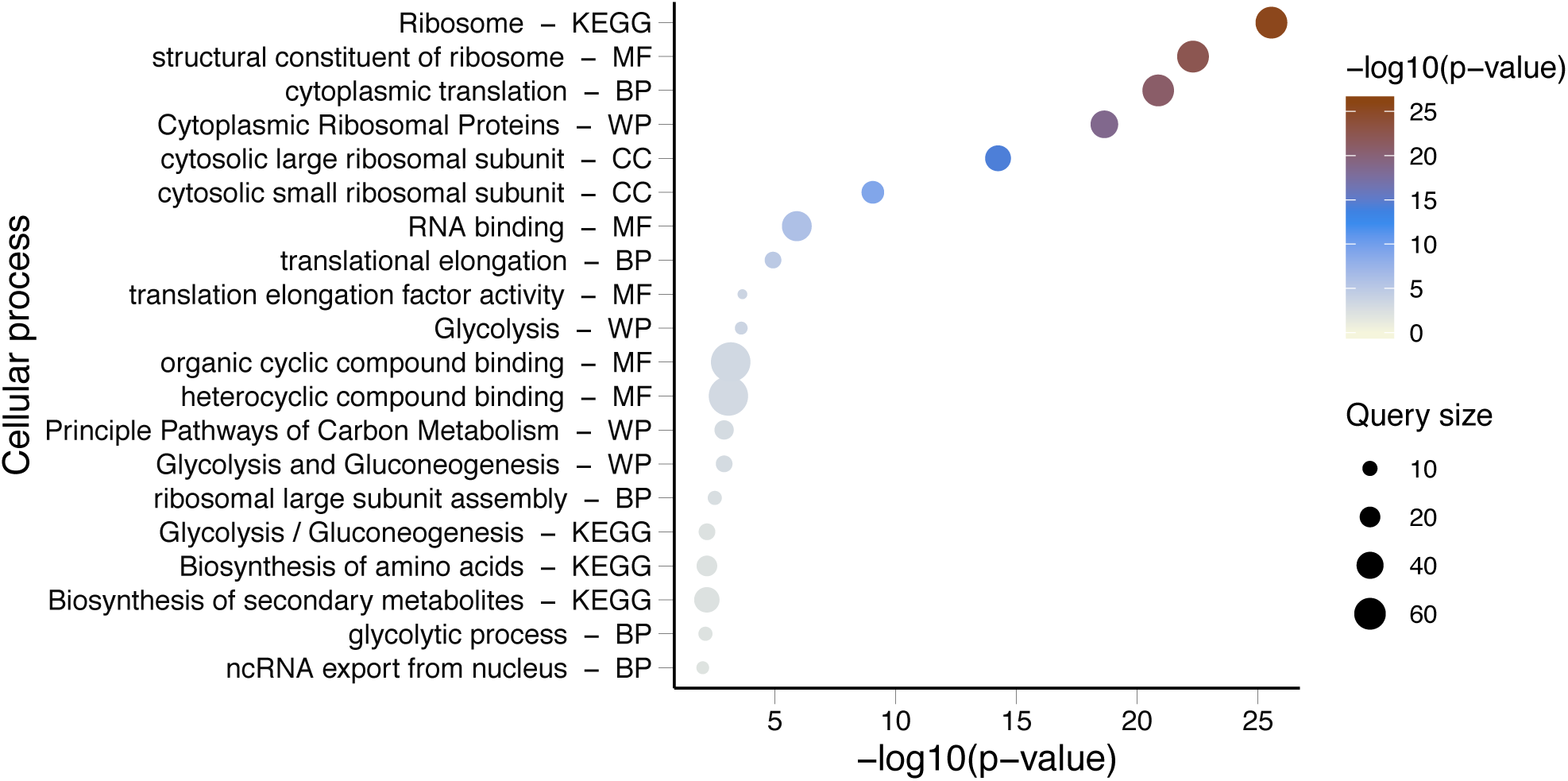
Hydrolysate-specific TF target gene functions. GO enrichment of TF target genes determined from ChIP-chip (Gonçalves et al., 2017) of TFs which modulate hydrolysate growth, generated using the gProfiler2 R package (Reimand et al., 2019).

**Supplementary Figure S7.**
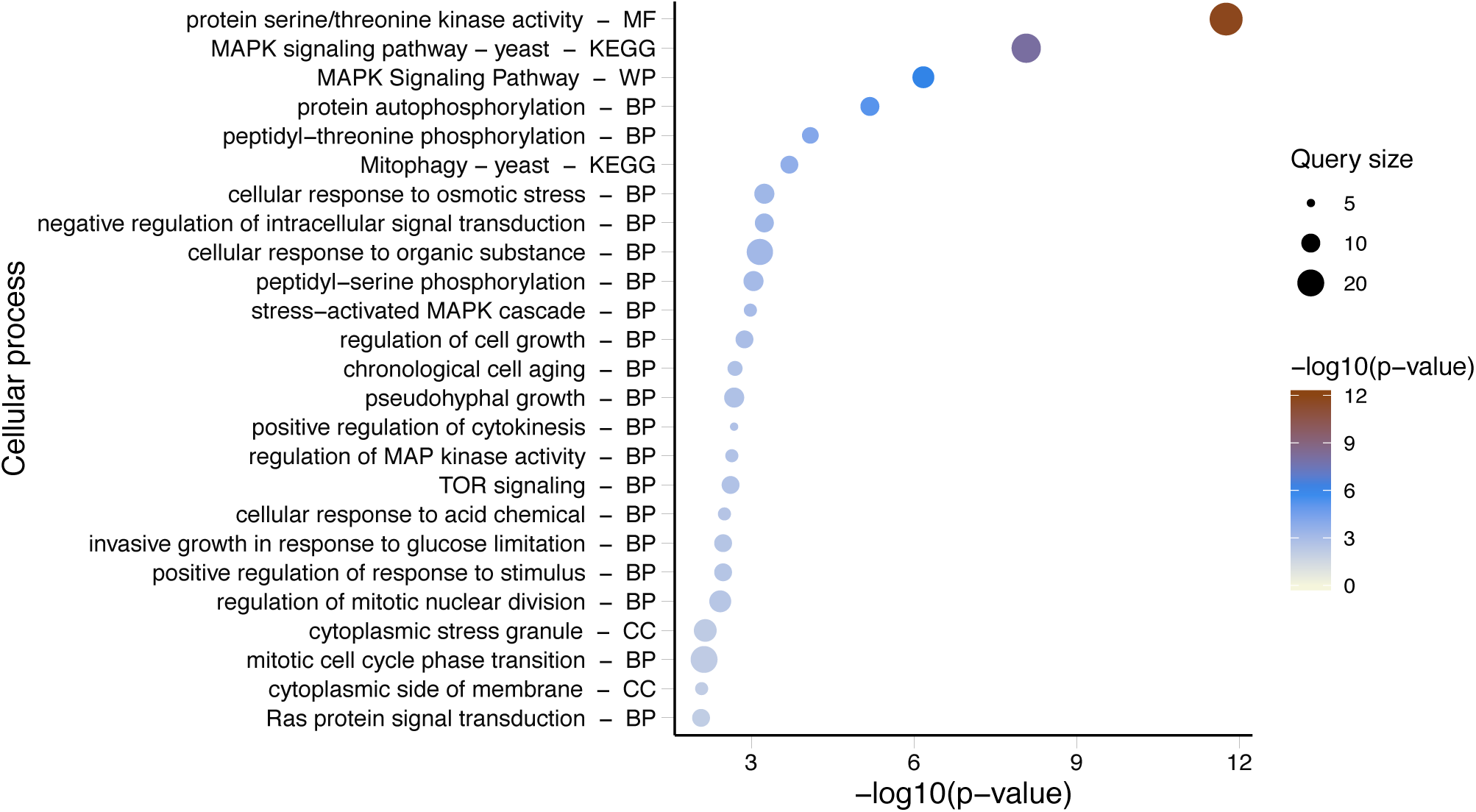
Hydrolysate-specific PK interactor functions. GO enrichment of PK phosphorylation targets determined from Phospho-proteomics data (Sadowski et al., 2013) of PKs which modulate hydrolysate growth, generated using the gProfiler2 R package (Reimand et al., 2019).

**Supplementary Figure S8.**
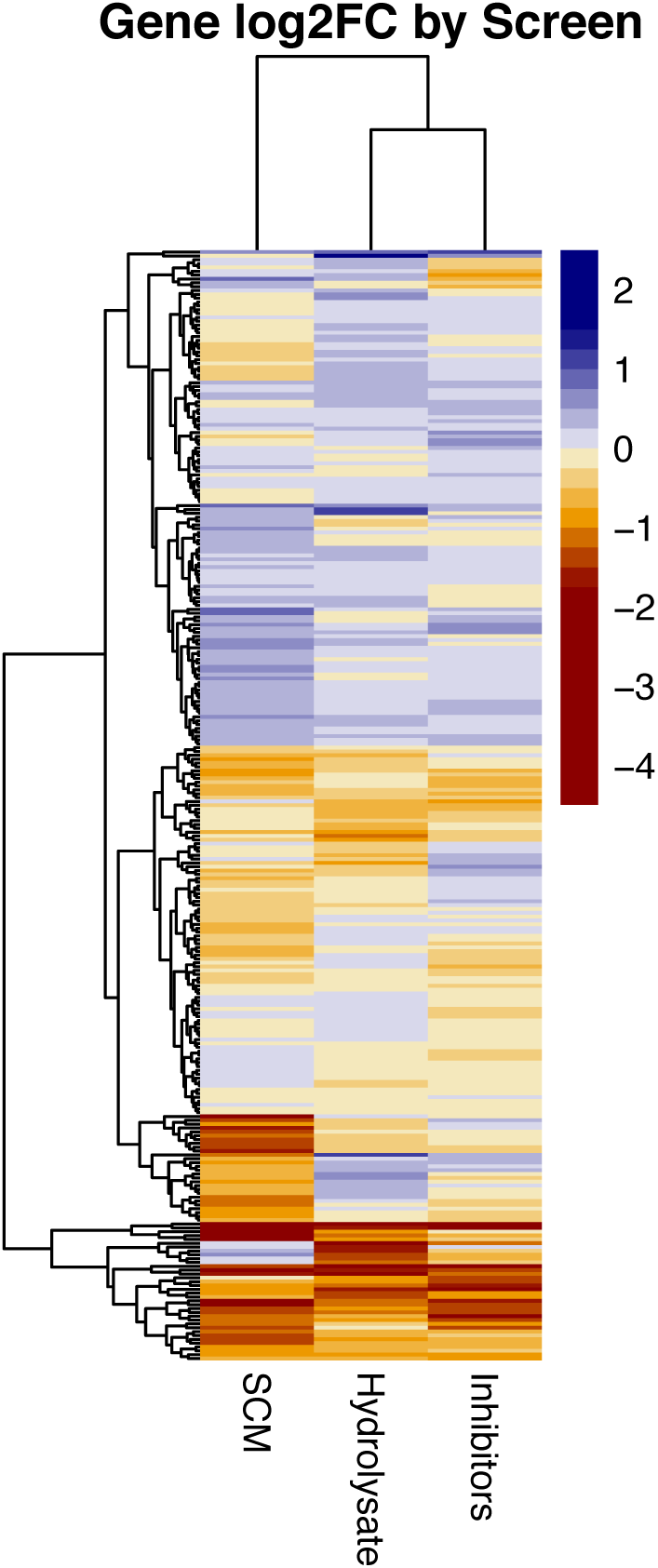
CRISPRi effects across screens. Log2 gene fold changes compared between SC medium, SCM + 10% Hydrolysate and SCM + 45% Inhibitor Cocktail. The heatmap was generated with the pheatmap R package (Kolde, 2019).

**Supplementary Table T1.** HPLC measurements. Metabolites concentrations (g l-1), measured by HPLC, of yeast cultures grown in SCM, SCM+10% hydrolysate and SCM+45% inhibitor cocktail. Glucose, ethanol and acetic acid concentrations were measured in the growth medium (Initial) and at the end of fermentations (Final). Three biological triplicates were performed for the three tested conditions, error represents the standard deviation between replicates.

|  | SCM |  | 10% Hydrolysate |  | 45 % Inhibitor cocktail |  |
| --- | --- | --- | --- | --- | --- | --- |
|  | Initial | Final | Initial | Final | Initial | Final |
| Acetic acid (g l<sup>-1</sup>) | 0.00 | 0.10 ± 0.03 | 0.62 | 0.35 ± 0.03 | 1.81 | 0.86 ± 0.13 |
| Ethanol (g l<sup>-1</sup>) | 0.00 | 4.32 ± 0.73 | 0.00 | 9.09 ± 0.76 | 0.00 | 4.32 ± 0.86 |
| Glucose (g l<sup>-1</sup>) | 20.06 | 0.00 ± 0.00 | 25.84 | 0.19 ± 0.17 | 20.03 | 0.00 ± 0.00 |

**Supplementary Table T2.**
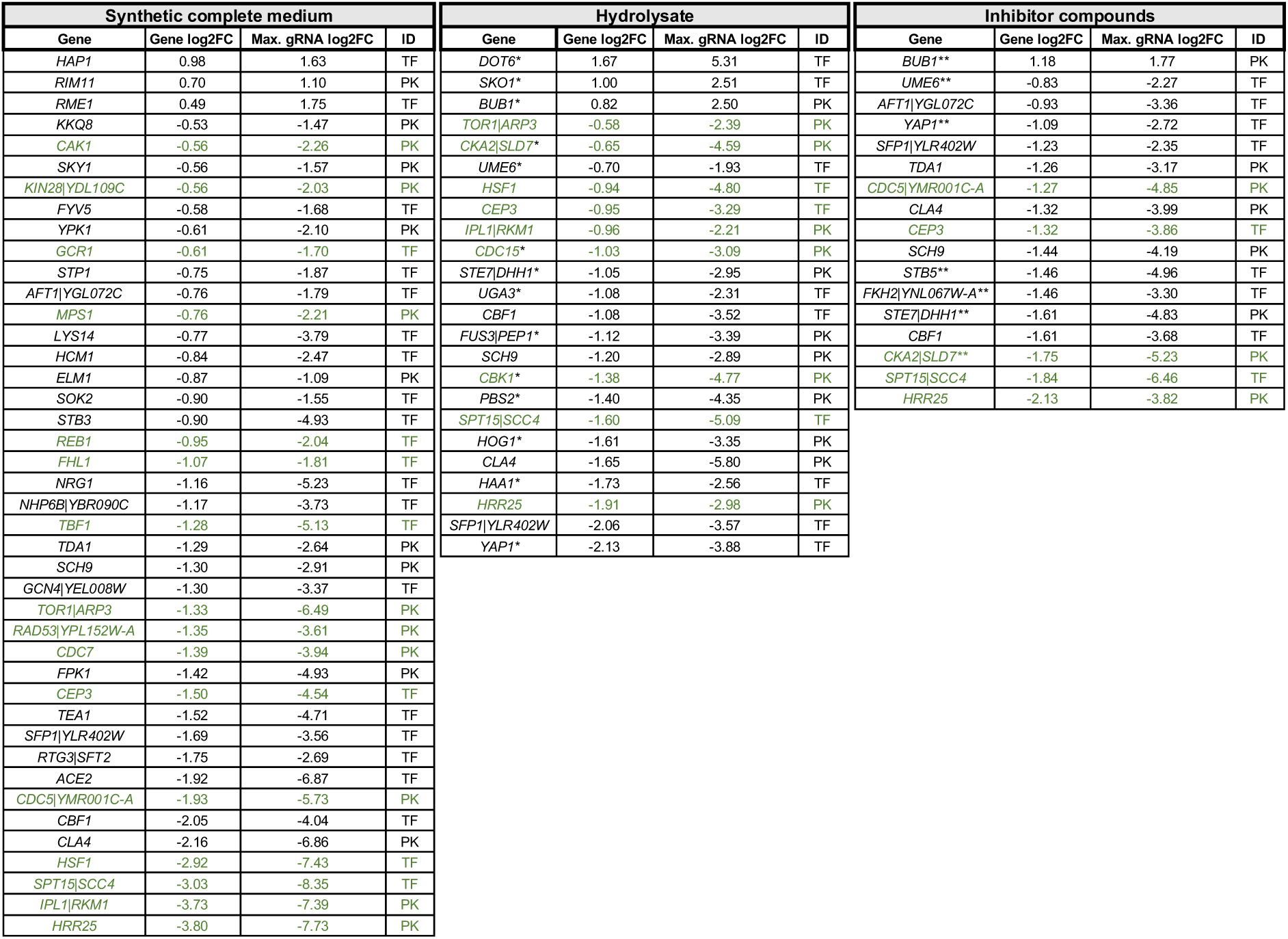
CRISPRi effects across media. Significant genes (gene level fold change with FDR<0.05 and at least two gRNAs with absolute log2FC>=1 and FDR<0.05) are shown across screens with their mean log2FC, their maximum gRNA log2FC and ID to specify if a TF or PK is targeted. For target genes transcribed from bidirectional promoters, both genes are reported (separated with a vertical dash). Tables are ordered by gene log2FC. Rows of essential genes as defined by in-viable knock-out mutants (Cherry et al., 1998) are in green color. One and two asterisks (*, **) behind a gene name indicate that repression caused hydrolysate-specific or inhibitor-specific effects, respectively (not measured in SCM).

